# NanoMOFs with Encapsulated Photosensitizer: Accumulation in *Chlamydia trachomatis* Inclusions and Antimicrobial Effects

**DOI:** 10.1101/2023.04.04.535539

**Authors:** Xiaoli Qi, Ekaterina Grafskaia, Zhihao Yu, Ningfei Shen, Elena Fedina, Alexander Masyutin, Maria Erokhina, Mathilde Lepoitevin, Vassili Lazarev, Nailya Zigangirova, Christian Serre, Mikhail Durymanov

## Abstract

Metal-organic framework nanoparticles (nanoMOFs) are a promising class of hybrid nanomaterials for biomedical applications. Some of them, including biodegradable porous iron carboxylates are proposed for encapsulation and delivery of antibiotics. Due to the high drug loading capacity and fast internalization kinetics nanoMOFs are more beneficial for the treatment of intracellular bacterial infections compared to free antibacterial drugs, which poorly accumulate inside the cells because of the inability to cross membrane barriers or have low intracellular retention. However, nanoparticle internalization does not ensure their accumulation in the cell compartment that shelters a pathogen. This study shows the availability of MIL-100(Fe) MOF nanoparticles to co-localize with *Chlamydia trachomatis*, an obligate intracellular bacterium, in the infected RAW264.7 macrophages. Furthermore, nanoMOFs loaded with photosensitizer methylene blue (MB) exhibit complete photodynamic inactivation of *Chlamydia trachomatis* growth. Simultaneous infection and treatment of RAW264.7 cells with empty nanoMOFs resulted in a 3-fold decrease in bacterial load that indicates an intrinsic anti-chlamydial effect of this iron-containing nanomaterial. Thus, our findings suggest the use of iron-based nanoMOFs as a promising drug delivery platform, which contributes to antibacterial effect, for the treatment of chlamydial infections.

## INTRODUCTION

The treatment of intracellular bacterial infections, such as *Chlamydiae*, is still challenging because the invasion of these pathogens to the host cells provides their protection from the host immune system and antibacterial drugs. Two membrane barriers, including the plasma membrane and the endosomal/phagosomal membrane of the host cell, significantly restrict access of antibiotics to intracellular bacteria.^1–4^ The developmental cycle of the obligate intracellular parasite *Chlamydia (C*.*) trachomatis* involves two alternating morphological forms such as the infectious elementary body (EB) and the proliferating reticulate body (RB). Upon internalization, EBs transform into metabolically active RBs, which modify a membrane of the intracellular vesicular compartment, termed “inclusion”, with bacterial proteins that prevent lysosomal fusion.^5^ The remodeled membrane subsequently provides migration of the inclusion towards the microtubule-organizing center (MTOC), which is in close proximity to the nutrient-rich peri-Golgi region.^6^ The major problem for the treatment of *C. trachomatis* infection is the ability of this parasite to transform into a metabolically inactive persistent form, called “aberrant RB”, upon the treatment with antibiotics (penicillin), gamma-interferon (IFNγ), or essential nutrient deprivation. However, *Chlamydia* restarts proliferation and dissemination after elimination of stressful stimuli. ^7^ Therefore, development of novel antibacterial drugs and antimicrobial drug delivery platforms is crucial for circumventing the defense mechanisms of *C. trachomatis*.

The use of nanoparticles as drug delivery systems for the treatment of intracellular bacterial infections is an attractive strategy.^2,3^ It has been shown that encapsulation of small-molecule antibacterial agents into nanocarriers can potentiate their efficacy. The benefits of nanoparticulate drug delivery systems (DDS) have been shown for the treatment of different intracellular infections including *Listeria monocytogenes, Mycobacterium spp*., *Staphylococcus aureus*, and others.^2,8^ Noteworthy, some efforts were made to develop nanoparticulate DDS for the treatment of chlamydial infections. For instance, Toti *et al*. demonstrated therapeutic effect of PLGA nanoparticles loaded with rifampin and azithromycin in *Chlamydia*-infected Hep-2 cells.^9^ Another group developed PAMAM dendrimer-azithromycin conjugates, which also displayed a strong antibacterial effect on the same cell model.^10^ Both mentioned nanoparticle drug delivery systems accumulated in chlamydial inclusions that probably enhanced the effectiveness of the antibiotics in reducing microbial burden. Thus, the ability of drug-loaded nanomaterials to concentrate in chlamydial inclusions seems to be a primary requirement for a potent antibacterial effect.

Here, we evaluated feasibility of using nanoparticles of biodegradable metal-organic framework (nanoMOFs) as drug delivery carriers for treatment of chlamydial infections. MOFs are a sub-class of coordination polymers formed by the self-assembly of metal ions and organic polydentate ligands.^11,12^ MOFs exhibit a large diversity of structures, including a large pore size, that when combined with their tunable and hybrid character, is suitable to encapsulate a large variety of therapeutic molecules with high efficiency and in most cases associated with a prolonged release profile.^13,14^ In addition, MOFs are biodegradable and can be used as a safe medication delivery vehicle that utilizes biocompatible compositions. Among them, the mesoporous iron (III) trimesate MOF denoted MIL-100(Fe) (MIL stands for “Materials of Institute Lavoisier”) is particularly attractive. Its architecture is constructed from iron oxoclusters connected by trimesate moieties resulting in a mesoporous cubic structure with two distinct cages. Its synthesis at the nanoscale under green ambient pressure conditions,^15^ its good biocompatibility and biodegradability in body fluids,^16^ as well as its ability to undergo surface functionalization,^17–19^ make this nanoMOF an appealing delivery platform for different small-molecule drugs.^20^

Although a few investigations have been conducted to evaluate the potential of MOFs as drug delivery systems for the treatment of bacterial infections,^21–25^ no studies focusing on *Chlamydiae* infections have been reported yet. To provide the rationale for using MIL-100(Fe) nanoparticles as a medication delivery vehicle to treat chlamydial infections, we evaluated here the internalization kinetics of these nanoMOFs and their potential to reach chlamydial inclusions in infected macrophages. Furthermore, the intrinsic antimicrobial activity of these materials and the feasibility of their use for reducing of the *Chlamydia* burden in infected cells was established. Finally, we produced these iron(III)-based nanoMOFs loaded with photosensitizer methylene blue (MB) for photodynamic treatment (PDT) of chlamydial infections (**Figure 1**). Once some of mucosal membranes infected by *C. trachomatis* are available for light treatment, PDT seems a feasible approach for eradication of this pathogen. Antibacterial photodynamic effect of MIL-100(Fe) nanoMOFs with encapsulated MB was evaluated in infected macrophages in comparison with free photosensitizer.

**Figure 1.**
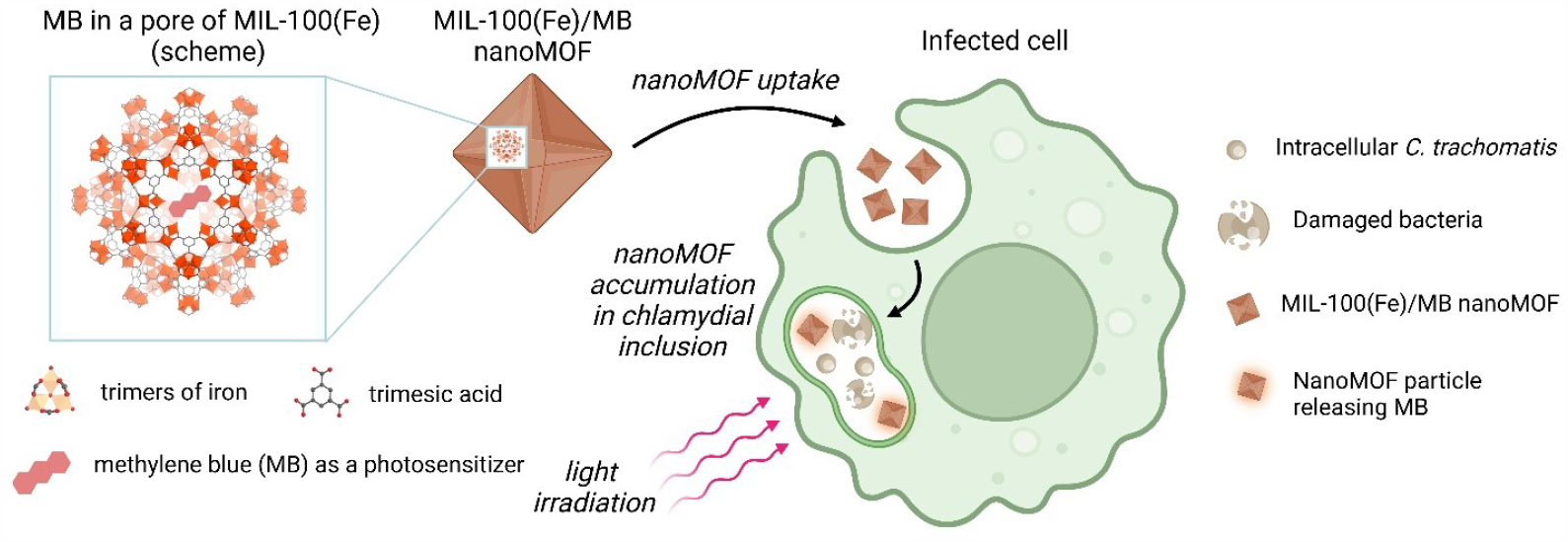
The scheme of using MIL-100(Fe)/MB nanoMOFs for photodynamic treatment of chlamydial infections.

## RESULTS AND DISCUSSION

### Characterization and Post-synthesis Modification of NanoMOFs

The physicochemical properties of nanoMOFs were analyzed by different methods (**Figure 2**). In deionized water, nanoMOFs exhibited a small hydrodynamic diameter, narrow size distribution, and a negative surface charge (**Table S1**). SEM microscopy (**Figure 2A,B**) showed that synthesized nanoparticles are significantly smaller than 100 nm in diameter. Due to their small size, nanoMOFs seem to be a highly efficient vehicle for drug delivery to intracellular compartments because of rapid internalization by different cell types *via* multiple endocytic pathways.^26^

**Figure 2.**
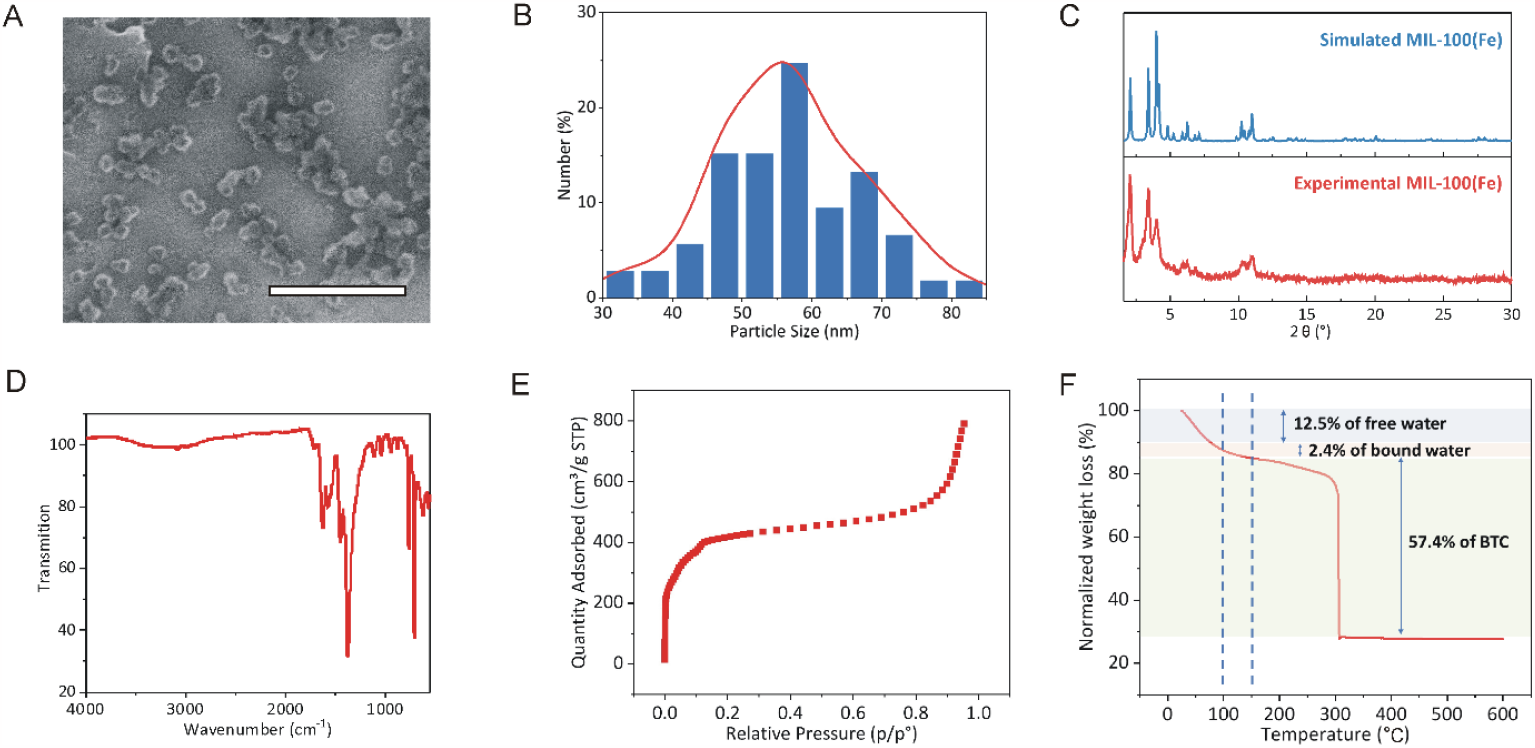
Physicochemical properties of MIL-100(Fe) nanoMOFs. (A) SEM image of nanoparticles. Scale bar is 500 nm. (B) Size distribution of nanoparticles determined by quantitative analysis of SEM images. Over 100 nanoparticles were measured. (C) Powder X-ray diffraction pattern of nanoparticles, which was recorded within 1.6°-30° 2θ range. (D) FT-IR spectrum of nanoMOFs within 4,000-550 cm^-1^ range. (E) Nitrogen adsorption isotherm of nanoparticles with BET surface area of 1,635(10) m^2^ per gram at 77 K. (F) Thermogravimetric analysis profile of nanoparticles under oxygen flow.

Using X-ray diffraction, the resulting peaks obtained at 2.06, 3.34, 4.06, 10.27 and 10.92° matched the simulated pattern for MIL-100 (Fe), which confirmed the structure of MIL-100(Fe) (**Figure 2C**). The slight broadening peak was the result of a small particle size (< 100 nm) (**Figure 2C**). FTIR spectrum (**Figure 2D**) shows the main characteristic peaks of MIL-100(Fe), which include the main characteristic carboxylate stretching vibration peaks at 1,670-1,220 cm^-1^, along with the peak at 620 cm^-1^ for stretching vibration of the Fe-µ3O bond. Except for the main peaks, there is a small peak at 1,702 cm^-1^, which is assigned to the stretching vibration of C=O groups due to residual traces of free trimesic acid.

N_2_ porosimetry at 77 K led to the expect sorption isotherm (**Figure 2E**) with the typical sub-steps characteristics of this bimodal mesopores solid, together with a calculated BET area of 1635 (10) m^2^ g^-1^.

TGA under oxygen atmosphere (**Figure 2F**) shows the typical weight losses from MIL-100(Fe) with the residue at high temperature attributed to Fe_2_O_3_. To assess the purity of the nanoMOFs, the mass of the dehydrated nanoparticles at 150°C was normalized as 100 %, showing a 64.6 wt% loss due to the degradation of the ligand at higher temperature, a value slightly higher than the theoretical one (63.2 wt%) most likely because of the residual traces of ligands trapped into the pores.

### Cellular Uptake and Intracellular Localization in RAW264.7 Macrophages

*C. trachomatis* is known to penetrate and propagate in different cell types that reflects the ability of this parasite to cause infections in different tissues. Cervical epithelium cells, fibroblasts and macrophages are the most used *in vitro* models for investigation of *C. trachomatis* biology.^27^ Here, we have investigated the ability of our nanoMOFs as a drug carrier to reach chlamydial inclusions in infected RAW264.7 macrophages as a cell model.

Nanoformulations as well as intracellular pathogens enter the macrophages by phagocytic route. For this reason, we evaluated the cellular uptake rate and intracellular accumulation of our nanoMOFs in non-infected RAW264.7 cells. The nanoparticles were fluorescently labeled with toluidine blue O (TBO) dye using a sulfo-NHS/EDC reaction. Flow cytometry analysis has shown that the internalization of nanoparticles in RAW264.7 cells rapidly increased in the first hour and reached a plateau after 4-6 hours of incubation (**Figure 3A,B**). More than 90 % of RAW264.7 cells were involved in the nanoparticle uptake after 1 hour of incubation. This is supported by a previous internalization report using these nanoMOFs.^26^ Most probably, the reduction in fluorescence intensity per cell and the fractions of cells containing TBO-labeled nanoMOFs after 6 hours is due to cell proliferation, because RAW264.7 cells are rapidly dividing cells with doubling time of 11 hours.^28^ To elucidate the fate of internalized nanoMOFs in cells, we studied their intracellular distribution by confocal microscopy. As expected, TBO-labeled nanoMOFs were co-localized with acidic vesicles inside the cells, and stained with a lysosome marker LysoTracker Green DND-26 dye (**Figure 3C**). This fits well with our previous findings that evidenced that MOF nanoparticles go mainly into the lysosomes after internalization.^26,29^

**Figure 3.**
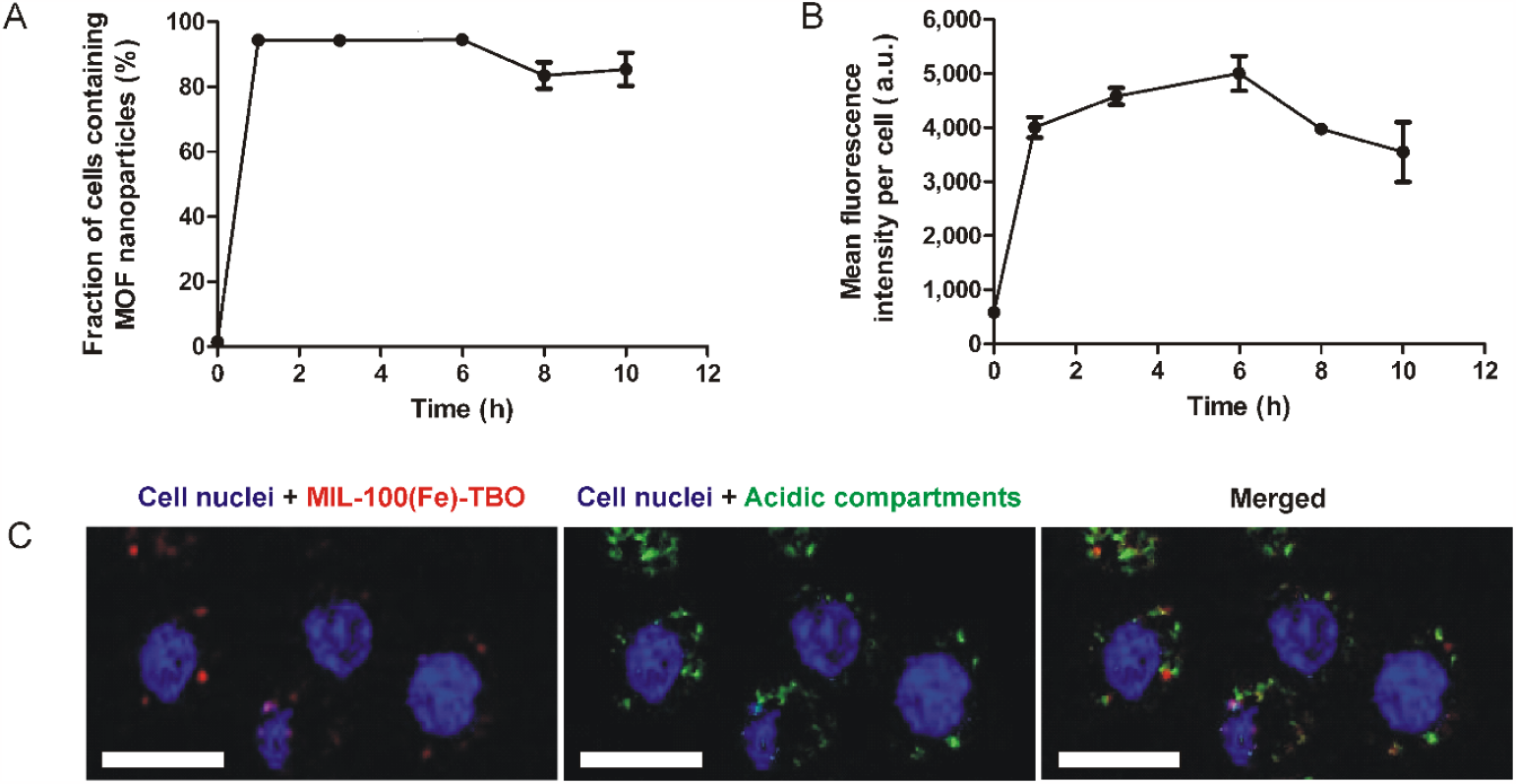
Internalization and intracellular localization of TBO-labeled nanoMOFs in RAW264.7 macrophages. (A) Cellular uptake kinetics of MIL-100(Fe)-TBO nanoparticles by RAW264.7 cells in terms of the fraction of cells containing fluorescently labeled nanoMOFs and (B) value of mean fluorescence intensity per cell. The data are given as means ± SD. (C) Intracellular localization of TBO-labeled nanoparticles in RAW264.7 cells. The incubation of cells with the fluorescently labeled nanoMOFs (red) after 5 hours resulted in their accumulation in acidic compartments stained with LysoTracker Green (green). Staining with Hoechst 33342 determines the nuclei of RAW264.7 cells (blue). Scale bar is 20 µm.

### Characterization of the Cell Model for Analysis of NanoMOF Interaction with Intracellular *C. trachomatis*

To establish a macrophage cell model infected with *C. trachomatis*, we aimed to find an optimal infection dose. It has been revealed that RAW264.7 cells are relatively resistant to *C. trachomatis* infection. According to obtained data, only a MOI of 100 resulted in a significant number of infected cells (**Figure S1**). This dose was selected for further experiments.

### NanoMOF Co-localization with *C. trachomatis* Inclusions in Infected Macrophages

The TEM-EDX technique was implemented to investigate intracellular interaction between MIL-100(Fe) nanoparticles and chlamydiae. It was found that chlamydiae accumulated in large vesicular compartments of infected macrophages (**Figure 4A**). In contrast, nanoMOFs in non-infected macrophages accumulated in relatively small phagosomes (**Figure 4B**). It was found that MOF nanoparticles can reach chlamydial inclusions of the infected RAW264.7 cells (**Figure 4C**). EDX analysis of the images indicated the presence of iron in co-localizing regions with bacteria particles, indicating them as nanoMOFs (**Figure 4D**). Most probably, it occurs due to MIL-100(Fe) nanoparticle accumulation in late endocytic vesicles, called multivesicular bodies (MVBs), followed by their direct fusion with chlamydial inclusions. It has been found in earlier literature that chlamydial inclusions can fuse with MVBs^30^ because *Chlamydia* uses MVBs as an additional nutrient source.^5^ Thus, the ability of nanoMOFs to accumulate in chlamydial inclusions can be further exploited for the delivery of different antibacterial drugs including antibiotics and photosensitizers.

**Figure 4.**
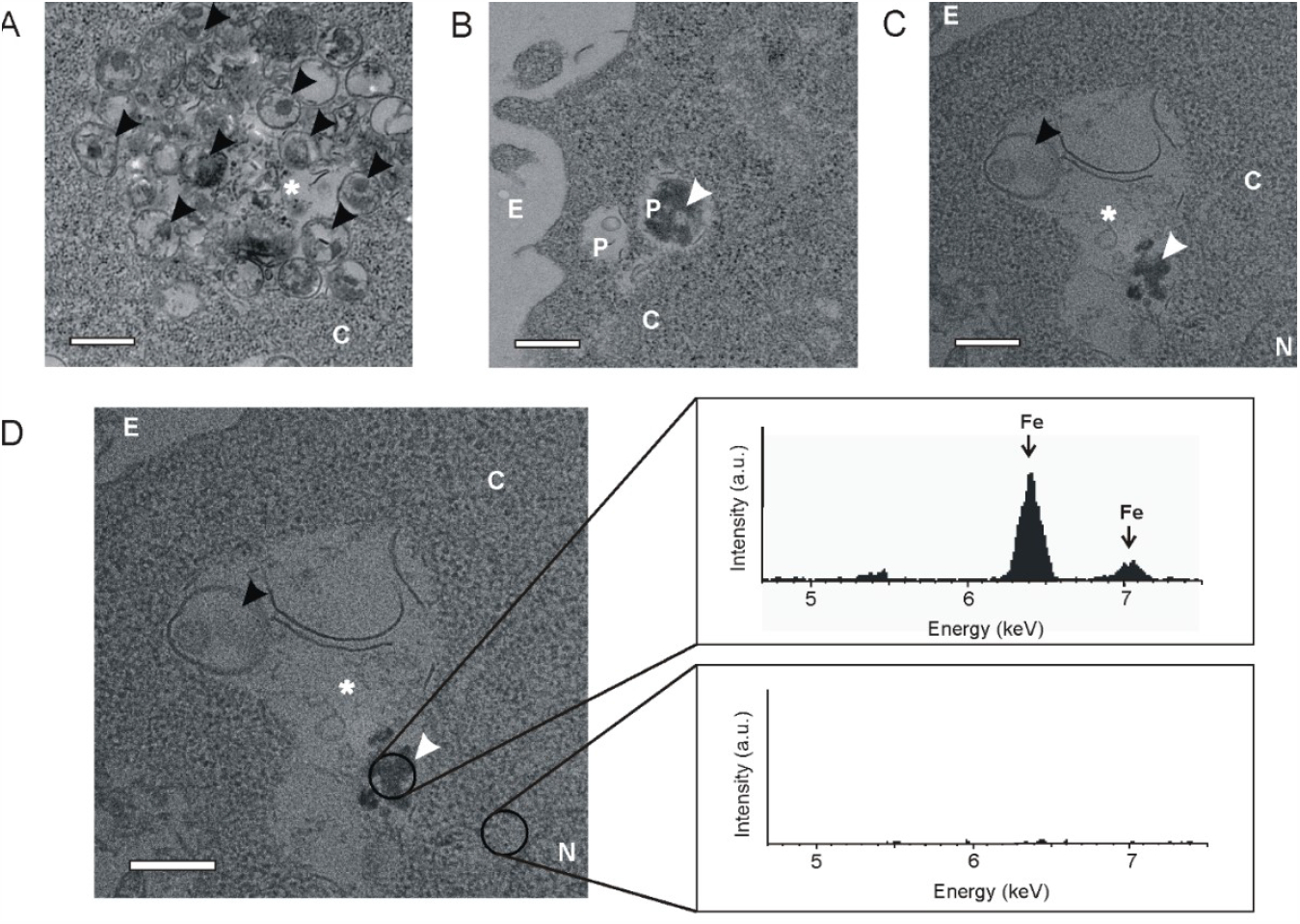
Analysis of MIL-100(Fe) nanoparticle accumulation in chlamydial inclusions using TEM imaging. (A) An image of chlamydial inclusion in cytoplasm of RAW264.7 cells infected with *C. trachomatis*. (B) Photograph of phagosome with nanoMOFs. (C) Co-localization of MIL-100(Fe) nanoparticles with *C. trachomatis* EBs within inclusions in RAW264.7 cells. (D) EDX spectrum analysis showing two major peaks of iron at 6.5 and 7.1 keV from the area with nanoMOFs. The peaks are not detected from the area without nanoparticles. (A)-(D) Black and white arrowheads indicate chlamydial EBs and MIL-100(Fe) nanoparticles, respectively. White stars indicate chlamydial inclusions. Abbreviations: N, nucleus; C, cytosol; P, phagosome; E, extracellular space. Scale bar is 500 nm.

### Intrinsic Antibacterial Activity of NanoMOFs against *C. trachomatis*

Noteworthy, previous studies have reported bactericidal effects of several MOF structures due to release of active metal cations or ligands.^31^ In our study, analysis of TEM images indicated damaged bacteria in most of the inclusions, where MOFs were present (**Figure S2**) that might indicate their antimicrobial properties. Intrinsic iron-based nanoMOF toxicity against *C. trachomatis* might also be a result of iron involvement into Haber-Weiss cycle. In fact, in macrophages this reaction becomes possible due to phagosome-associated NADPH oxidase, which generates O_2_^·-^ in the phagosomal lumen.^32^ As a result, iron from nanoMOFs could accelerate conversion of O_2_^**·-**^ to ^·^OH that would lead to bactericidal effect.

It turned out that the addition of MIL-100(Fe) nanoparticles to RAW264.7 cells at the time of infection resulted in a significant reduction in size of chlamydial inclusions (**Figure 5A,B**). We further quantified the infectious chlamydial progeny recovered from the treated RAW264.7 cells and transferred to HeLa cells (**Figure 5C,D**). A remarkable 3-fold decrease in *Chlamydia* titer obtained from nanoMOF-treated RAW264.7 cells was observed (**Figure 5E,F**). Intrinsic bactericidal activity of these iron (III) trimesate nanoMOFs could be advantageous in terms of antibacterial drug delivery once it contributes to therapeutic effect.

**Figure 5.**
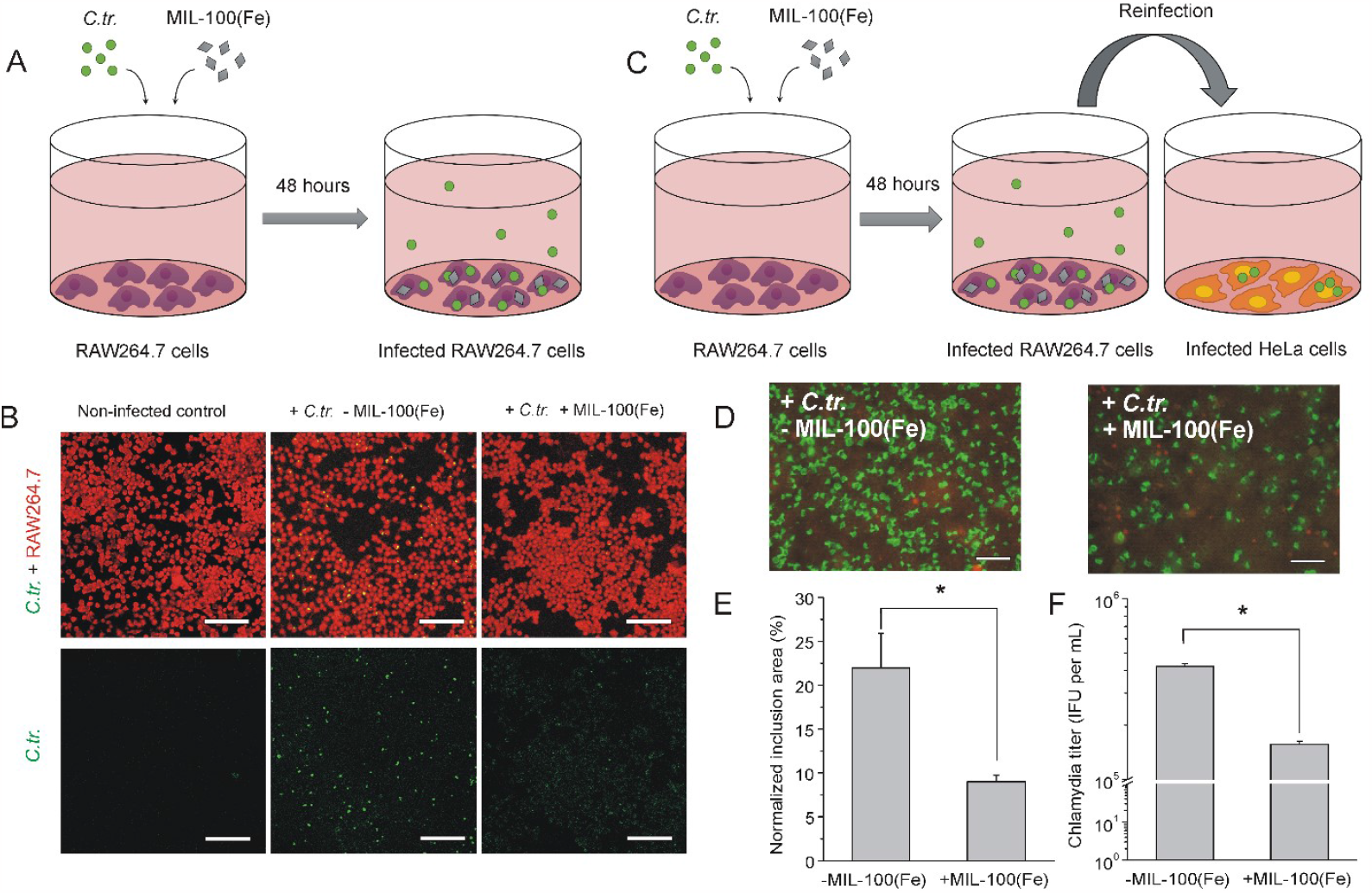
Intrinsic antibacterial effect of nanoMOFs. (A) The scheme of RAW264.7 cell infection and simultaneous treatment with MIL-100(Fe) nanoparticles. (B) After 48 hours of incubation, RAW264.7 macrophages were stained with Evans blue to visualize the cells (red) and FITC-labeled antibodies against chlamydial major outer membrane protein to visualize chlamydial inclusions (green). (C) The scheme of the experiment with reinfection of HeLa cells to determine infectivity of chlamydial progeny recovered from RAW264.7 macrophages, infected with *C. trachomatis* and treated or non-treated with nanoMOFs. (D) Images of reinfected HeLa cells (red) with chlamydial inclusions (green) visualized by staining with FITC-labeled antibodies against chlamydial major outer membrane protein. Scale bar is 50 µm for (B) and (D). (E) Quantitative analysis of reinfection efficacy determined as normalized chlamydial inclusion area. (F) Quantitative analysis of reinfection efficacy determined as infectious progeny yield. Data are shown as means ± SD. * p < 0.05, Mann-Whitney U test.

### Release Behavior and Photodynamic Activity of MB-loaded NanoMOFs *in vitro*

The ability of the iron (III) nanoMOFs to accumulate in chlamydial inclusions enables their use as a carrier for delivery of antibacterial drugs. As mentioned above, *Chlamydia* can transform to metabolically inactive persistent form and become resistant to the action of antibiotics, which interfere with bacteria metabolism. For this reason, we selected photosensitizer as an antimicrobial cargo, whose action does not rely on metabolism interference.

To fabricate nanoMOFs for photodynamic treatment, photosensitizer methylene blue (MB) was therefore encapsulated into MIL-100(Fe) nanoparticles with a loading extent of 2 wt%. Analysis of drug release behavior indicated a minor release of MB in milliQ (deionized) water. However, the resuspension of MB-loaded nanoMOFs in different physiological solvents led to a significant increase in the rate of MB release (**Figure 6A**). The most rapid release of MB was observed in PBS, resulting in almost 100 % release in 24 hours at 37⁰C. This is mainly due to the progressive decomposition of the nanoMOFs in the presence of phosphates. Our data are in compliance with the results of the recent study, where 30 % release of the organic linker (trimesate) from the nanoMOF was released upon 24 hours of incubation in PBS.^33^ It has been shown in the same study that the addition of bovine serum albumin could significantly slowdown nanoMOF degradation in PBS presumably due to the formation of protein corona around the particles and becoming a steric barrier for diffusion of phosphates inside the MOF structure.^33^ Indeed, we found that MB release in fetal bovine serum (FBS) occurred in a more sustained manner resulting in 80 % of released photosensitizer after 48 hours of incubation at 37⁰C. Probably, the slower release profile also could be explained by the delayed biodegradation of MIL-100(Fe) nanoparticles in FBS due to the opsonization of MOF nanoparticles with serum proteins.

**Figure 6.**
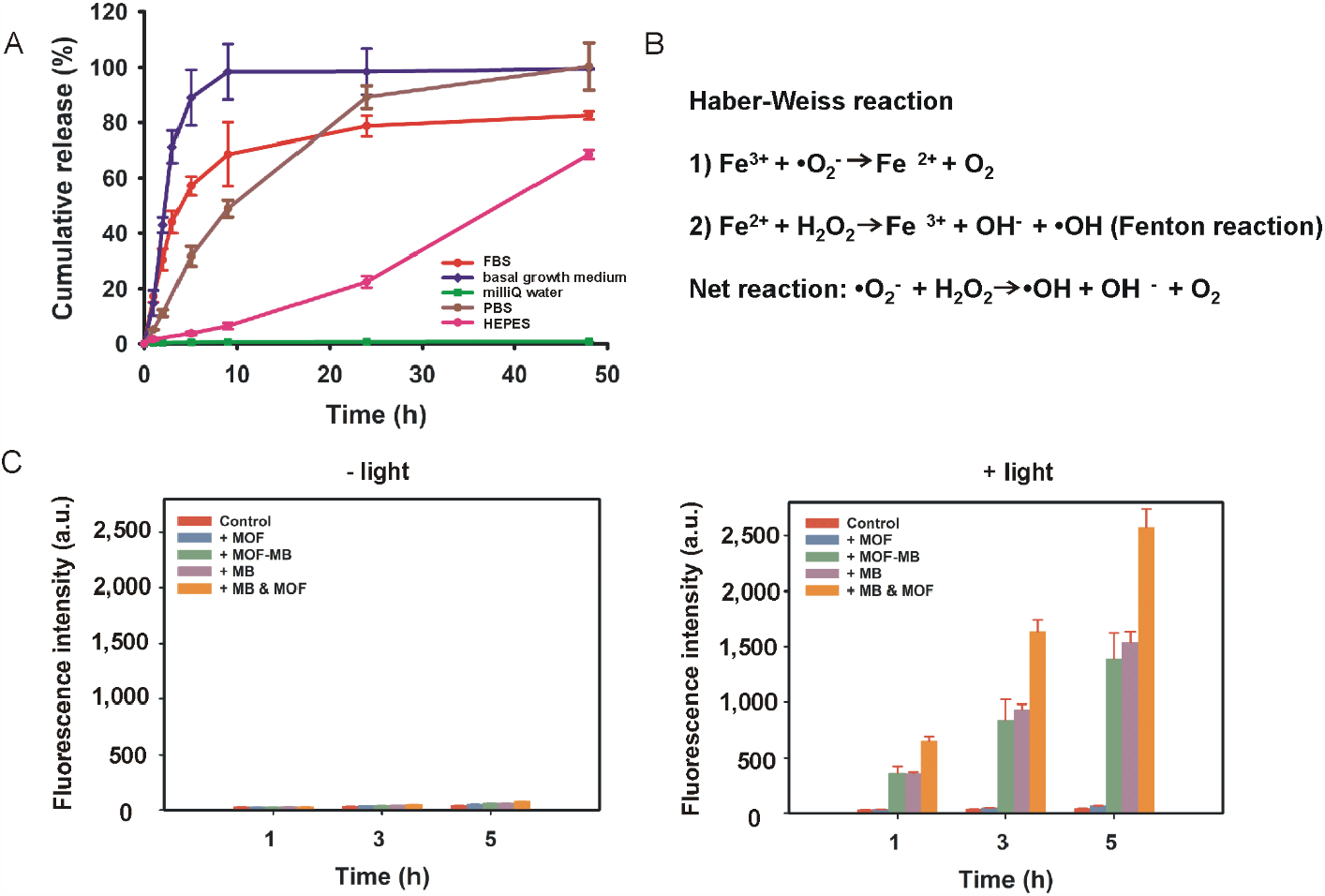
Release behavior and photodynamic activity of MB-loaded nanoMOFs *in vitro*. (A) Kinetics of MB release from MIL-100(Fe)/MB nanoparticles in different solutions within 48 hours of incubation at 37⁰C. (B) The scheme of Haber-Weiss reaction. (C) *In vitro* photodynamic effect of different formulations measured by fluorescence of H_2_DCF-DA probe. The photodynamic effect develops only in formulations, containing MB, under light irradiation. Control: buffer; MOF: nanoMOF; MOF-MB: MB loaded nanoMOFs; MB: free MB; MB&MOF: a mixture of free MB and nanoMOF. (A), (C) Data are shown as means ± SD.

We hypothesized that in terms of photodynamic therapy MIL-100(Fe) nanoparticles can potentiate the bactericidal effect of photosensitizer because of iron involvement in the Haber-Weiss reaction (**Figure 6B**). To measure ROS generation *in vitro*, we used H_2_DCF-DA acting as a ROS probe. Usually, this assay relies on the intracellular H_2_DCF-DA hydrolysis to H_2_DCF mediated by the cellular esterases, followed by a ROS-mediated oxidation of H_2_DCF to the fluorescent DCF.^34^ Our data suggest that the hydrolysis of H_2_DCF-DA can occur in a cell-free manner (without participation of the esterases) because we observed an increase in fluorescence in the light-treated samples and did not observe it the dark control. Time-dependent character of fluorescence increase in the MB-containing samples after a light exposure indicates a formation of oxidated non-fluorescent products, which become fluorescent upon hydrolysis over time.

This assay has shown that incubation of the nanoMOFs with free MB after light irradiation led to a 1.5-fold increase of ROS generation over time in comparison with free MB. Therefore, these MOF nanoparticles enhance the photodynamic effect of MB (**Figure 6C**). Surprisingly, the nanoMOF-encapsulated MB did not demonstrate any increase in ROS generation as compared with free MB after light exposure (**Figure 6C**). According to obtained UV-VIS spectra, it is due to quenching processes in MB-loaded MOFs (**Figure S3**). This highlights the significance of the MB release for MOF-potentiated photodynamic effect. Although *in vitro* release of photosensitizer from nanoMOFs occurs gradually and lasts depending on the solvent from 24 to 48 hours (**Figure 6A**), degradation of MOF structure inside the cells starts in 15-20 minutes after internalization as was recently shown on a macrophage model for MIL88B-NH_2_(Fe),^29^ another Fe(III) carboxylate MOF structure. It means that MIL-100(Fe)-mediated enhancement of photodynamic effect upon photosensitizer release is quite possible inside intracellular vesicular compartments.

### Photodynamic Effect of MIL-100(Fe)/MB Nanoparticles in Infected RAW264.7 Macrophages

An ideal antibacterial drug should provide complete eradication of intracellular parasite and should not be harmful to a host cell. A cytotoxicity analysis of nanoMOFs, with or without MB, has shown that they remained non-toxic towards non-infected RAW264.7 cells in a wide range of concentrations (**Figure S4A**). The concentration of MB-loaded nanoMOFs, which was used in this study for the treatment of infected macrophages, caused insignificant cytotoxicity at the selected dose of light irradiation (**Figure S4B**).

It turned out that the treatment of infected RAW264.7 cells with the MB-loaded nanoMOFs and other formulations, followed or not by light irradiation, did not significantly alter an appearance of chlamydial lipooligosaccharide in macrophages as was shown by ICC staining (**Figure S5**). Noteworthy, ICC staining does not reflect the parasite viability. Therefore, for testing the infectivity of the pathogen in the treated samples we carried out a reinfection assay using HeLa cells (**Figure 7A**).

**Figure 7.**
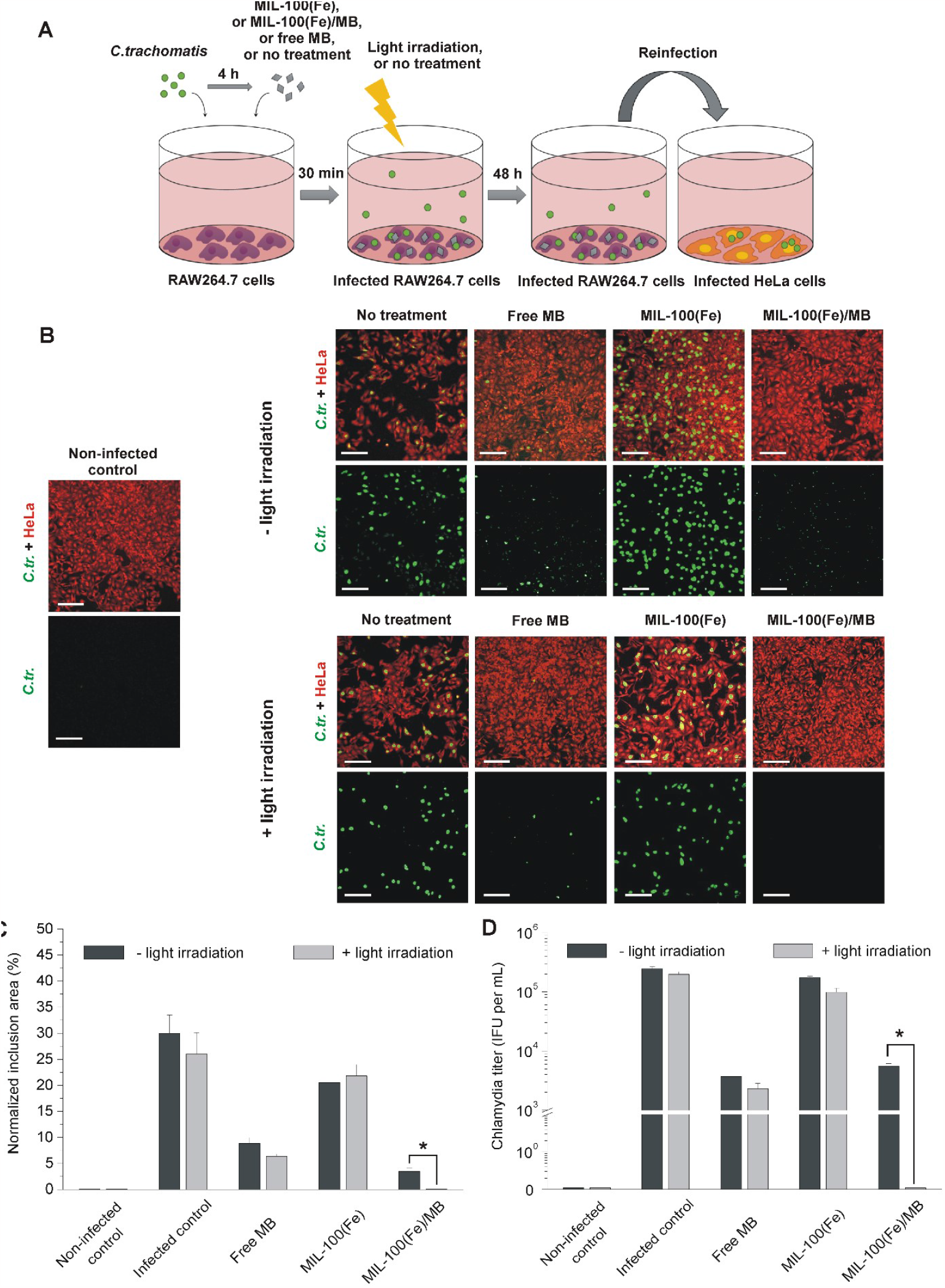
Photodynamic effect of MB-loaded nanoMOFs in *C. trachomatis*-infected RAW264.7 macrophages. (A) The scheme of the reinfection experiment with HeLa cells to determine an infectious progeny yield of infected RAW264.7 macrophages treated with different formulations, followed by light irradiation. (B) Images of reinfected HeLa cells (red) with chlamydial inclusions (green) visualized by staining with FITC-labeled antibodies against chlamydial lipooligosaccharide. Scale bar is 50 µm. (C) Quantitative analysis of the reinfection efficacy. To quantify an infection rate in HeLa cells, we normalized the area of inclusions to the area of HeLa cells. (D) Quantitative analysis of the reinfection efficacy determined as an infectious progeny yield. Data are shown as means ± SD. * p < 0.05, Mann-Whitney U test.

It was found that MB-loaded nanoMOFs significantly reduced infectivity titer even without light irradiation, while the bare nanoMOFs exhibited an insignificant decrease in infectious progeny yield (**Figure 7B-D**). Dark control with free MB treatment also led to substantial inhibition of *Chlamydia* growth upon reinfection of HeLa cells indicating anti-chlamydial effect of this dye. Interestingly, bactericidal effect of MB was reported earlier for different bacteria species in several studies,^35–37^ although its molecular mechanisms remain elusive. It should be noted that we did not detect here a significant reduction of *Chlamydia* titer in RAW264.7 cells treated with empty nanoMOFs (**Figure 7B-D**). Perhaps, such a lack of notable antibacterial effect could result from a much smaller nanoMOF accumulation in chlamydial inclusions as compared with treatment, where nanoMOFs were added during infection.

Light irradiation significantly enhanced therapeutic effect of MB-loaded nanoMOFs resulting in complete inhibition of chlamydial growth as compared with non-irradiated control. Meanwhile, no significant potentiation was observed for free MB and the MIL - 100(Fe) nanoMOFs after light treatment (**Figure 7B-D**). Eradication of *C. trachomatis* from infected RAW264.7 macrophages using MIL-100(Fe)/MB nanoMOFs could be attributed to photosensitizer delivery to chlamydial inclusions and MIL-100(Fe)-mediated enhancement of photodynamic effect presumably *via* the Haber-Weiss reaction as mentioned above. We used red light, from the visible spectrum, for the treatment of infected macrophages. It has been reported earlier that *C. trachomatis* is sensitive to water-filtered infrared irradiation by wavelength over 750 nm.^38,39^ Feasibility of this strategy was shown on a model of an eye infected with *Chlamydia*. It should be noted that a 5-fold decrease in infectious titer was achieved at a 4-folds higher irradiation dose than in the case of our approach. We used a dose rate of only 120 J cm^-2^, which did not inhibit *C. trachomatis* in absence of MB or MB-loaded nanoMOFs. Thus, our strategy provides delivery of photosensitizer to *C. trachomatis* inclusions and improves antibacterial photodynamic effect upon low-dose light exposure.

Thus, our results clearly demonstrate feasibility of photodynamic therapy for the treatment of chlamydial infection and improved efficacy of this treatment in a case of photosensitizer encapsulation into MIL-100(Fe) nanoparticles.

## CONCLUSION

In this study we have shown that MIL-100(Fe) nanoMOFs were able to accumulate in chlamydial inclusions. It means that application of these nanoparticles can be used as a drug delivery platform for different antibacterial agents that provides enhanced therapeutic effect against intracellular *C. trachomatis*, as shown for MB-loaded nanoMOFs. Second, these nanoparticles exhibit intrinsic bactericidal properties against this pathogen that can potentiate efficacy of delivered drugs.

Based on these findings, we first developed a photodynamic strategy for eradication of *Chlamydiae* that could find a potent application for the treatment of cervical or conjunctival infections. We also believe that the developed approach could be valuable for treatment of persistent chlamydial infections because it does not rely on inhibition of bacteria metabolism as compared with antibiotics.

## METHODS

### Synthesis and Characterization of MIL-100(Fe) Nanoparticles

MIL-100(Fe) nanoparticles (nanoMOFs) were synthesized through a green ambient pressure synthesis method reported by us.^15^ The nanoMOFs were characterized by means of powder X-ray diffraction (PXRD), thermogravimetric analysis (TGA), nitrogen porosimetry, Fourier-transform infrared (FTIR) spectrometry, dynamic light scattering (DLS), and scanning electron microscopy (SEM).

Scanning electron microscopy (SEM) images were obtained using FEI Magellan 400 scanning electron microscope. Zeta potential and dynamic light scattering (DLS) size measurement of hydrodynamic diameters were performed on a Malvern Zetasizer NanoZS (Malvern Instruments). Samples for measurements were prepared at final concentration of 0.1 mg mL^-1^. PXRD data were recorded using high-throughput Bruker D8 Advance diffractometer working in transmission mode and equipped with a focusing Göbel mirror producing CuKα radiation (λ_Cu_ = 1.5418 Å) and a LynxEye detector. Infrared spectra were measured with Nicolet iS5 FTIR ThermoFisher spectrometer in the 4000-500 cm^-1^ range. Nitrogen porosimetry data were collected using Micromeritics Tristar instrument at 77K (pre-activating samples at 150°C under vacuum for 8 hours). Thermogravimetric analysis (TGA) was performed using Mettler Toledo TGA/DSC 2, STAR System apparatus with a heating rate of 5°C per min under the oxygen flow.

### Fluorescent Labeling of MIL-100(Fe) Nanoparticles

Toluidine Blue O (TBO) dye was conjugated to the external surface of MIL-100(Fe) nanoparticles using EDC/NHS coupling reaction. NanoMOFs (5.0 mg) were dispersed in 1.2 mL of 5 mM HEPES buffer (pH 7.4) and sonicated for 2 minutes. Then, 1.6 mg EDC and 2.2 mg NHS were dissolved in 700 μL of HEPES and added to the nanoMOF suspension, followed by 30 min stirring at room temperature. Then, 100 μL of TBO (1 mg mL^-1^) in anhydrous DMSO was added to the mixture and stirred for two hours at room temperature. After the reaction, nanoMOFs were collected by centrifugation at 20,000×g for 20 min and washed 3-4 times with ethanol and water to remove unreacted TBO. TBO-labeled nanoMOFs were then lyophilized for 24 hours. The labeling ratio was determined by measuring OD of MIL-100(Fe)-TBO suspension upon decomposition, induced by 24-hour incubation in 200 mM citrate buffer (pH 5.5) at 40°C. OD was measured using CLARIOstar PLUS plate reader (BMG Labtech, Offenburg, Germany) at λ_max_ = 626 nm.

### Cell Culture

Murine RAW264.7 macrophages (ATCC, TIB-71) and human cervical adenocarcinoma HeLa cells (ATCC, CCL-2) were cultured in DMEM growth medium containing L-glutamine and pyruvate, 1 % PenStrep, 1 % nonessential amino acids, and 10 % fetal bovine serum. The cells were grown in a humidified incubator with 5 % CO_2_ at 37°С. The cells were used in experiments at passages 17-25 and 10-15 for RAW264.7 and HeLa cells, respectively.

### Flow Cytometry Analysis of MIL-100(Fe)-TBO Nanoparticle Uptake Kinetics by RAW264.7 Macrophages

RAW264.7 cells were seeded in 24-well plates (3,000 cells per well) 24 h before addition of MIL-100(Fe)-TBO nanoparticles at final concentration of 5 μg mL^− 1^. After 1, 3, 6, 8, and 10 h, growth medium was removed, and the cells were washed once with HBSS and resuspended in Versene solution. The internalization of nanoMOFs was analyzed using CytoFLEX (Beckman Coulter, Brea, CA) flow cytometer, equipped with 640 nm laser. At least 10,000 events per sample (3 samples per time point) were gated in 625-675 nm channel.

### Intracellular Localization of TBO-labeled NanoMOFs in RAW264.7 Macrophages

For analysis of intracellular localization of MIL-100(Fe)-TBO nanoparticles, RAW264.7 cells were seeded in 24-well plates and cultured for 24 h in an incubator at 37°С with 5% CO_2_ before addition of nanoMOFs at final concentration of 5 μg mL^-1^ and followed by 3-hour incubation. Before imaging, the cells were treated with LysoTracker Green DND-26 and Hoechst 33342 for labeling of acidic compartments and cell nuclei, respectively. The cell images were acquired in multitrack mode using confocal microscope Zeiss LSM 710 (Carl Zeiss, Oberkochen, Germany), equipped with Plan-Apochromat ×63/1.4 Oil DIC lens.

### *C. trachomatis* Culture and Infection

*C. trachomatis* strain L2/434/Bu (ATCC VR-902B) was routinely propagated in McCoy cells (ATCC CRL-1696) and isolated by urografin gradient ultracentrifugation as described previously.^40^ Isolated EBs were resuspended in a sucrose/phosphate/glutamic acid buffer (SPG) and stored at -70°C. *C. trachomatis* titers were determined by infecting cell monolayers with decimal dilutions of the stock suspension.

### Analysis of *C. trachomatis* and NanoMOF Co-localization in RAW264.7 Macrophages

To investigate intracellular co-localization of chlamydial EBs and nanoMOFs, RAW264.7 cells were seeded in 24-well plate at a density of 20,000 cells per well and incubated overnight. *C. trachomatis* L2/434/Bu at a MOI of 100 were added, followed by centrifugation for 2 hours at 1,500 rpm and incubation at 37°C and 5 % CO_2_ atmosphere for 2 hours. Then, the cells were washed with PBS and nanoMOFs were added at final concentration of 100 µg mL^-1^, followed by incubation for additional 2 hours. The samples with treated RAW264.7 cells were washed with 0.1M HEPES buffer (pH 7.4) and fixed with 2.5 % of glutaraldehyde (Ted Pella, Inc) in the same buffer for 2 hours. Then, the cells were washed from the fixative and postfixed with a 1 % solution of osmium tetroxide (OsO_4_) in 0.1 M HEPES for 1 hour at +4℃. Next, the samples were dehydrated by a graded series of ethanol and acetone, followed by embedding into epoxy resin. Ultrathin sections (60 -80 nm) were obtained using Leica UCT 4 ultramicrotome (Leica, Wetzlar, Germany) and stained with uranyl acetate. Obtained samples were analyzed using JEM 2100 microscope (Jeol, Tokyo, Japan). EDX analysis was performed using X-Max 8 mm^2^ detector (Oxford Instruments, Abingdon, UK) connected to a vacuum chamber of an analytical transmission electron microscope (JEM 2100, Tokyo, Japan) with Inca software (Oxford Instruments, Abingdon, UK).

### Analysis of MIL-100 (Fe) Bactericidal Effects

Antibacterial effect of nanoMOFs was examined on RAW264.7 cells infected with *C. trachomatis* at a MOI of 100. The nanoparticles were added to RAW264.7 cells during infection at a final concentration of 100 µg mL^-1^. After 48 hours of incubation, RAW264.7 cells were fixed with methanol and stained with Evans Blue for cell visualization. Chlamydial inclusions were visualized by staining with Pathfinder *C. trachomatis* Direct Specimen System (#30704, BIO-Rad) according to manufacturer’s recommendations. Another 96-well plate with RAW264.7 cells was frozen at -70⁰C to obtain cell lysates. RAW264.7 cell lysates were diluted in IMDM growth medium and transferred to the wells of 24-well plate, containing HeLa cell monolayer. Upon 48 hours of incubation, Hela cells were also immunocytochemically (ICC) stained in the same manner. Then, stained RAW264.7 and HeLa cells were photographed using the confocal microscope Zeiss LSM 710 (Carl Zeiss, Oberkochen, Germany) equipped with a Plan 10x/0.25 objective. The number of HeLa cells, containing chlamydial inclusions, was determined by calculation of positive cells in 30 random fields. The infection rate was also determined as the area of inclusions normalized to the area occupied by HeLa cells. The images of the cells were quantified using Fiji software (version 2.1.0/1.53c).

### Generation and Characterization of MB-loaded NanoMOFs

Encapsulation of methylene blue (MB) into nanoMOFs was performed by a simple impregnation method. First, the nanoparticles (3.5 mg) were dispersed in 1.33 mL of Milli-Q (deionized) water and sonicated for 2 minutes at room temperature. Next, 70 μL of MB (1 mg mL^-1^) in Milli-Q water was added to MOF suspension, followed by 24 h stirring at room temperature. Then, MIL-100(Fe)/MB nanoMOFs were collected by centrifugation (10 min, 17,000×g, room temperature), washed three times with Milli-Q water to remove free MB, collected again, and freeze-dried. The loading extent (wt%) was determined by measurement of MB optical density at λ = 660 nm after decomposition of MB loaded nanoMOFs by 24-hour incubation in 200 mM citrate buffer (pH 5.5) at 40°C.

The release profiles of MB from MB-loaded nanoMOFs were determined in different buffer solutions including Milli-Q, HEPES, PBS, fetal bovine serum (FBS) and StemPro® MSC SEM basal growth medium. Briefly, MB-loaded nanoparticles were dispersed in 1 mL of buffer solution and kept in the incubator at 37°C with mild shaking. At different incubation time points (1, 3, 5, 9, 24, and 48 h), 0.5 mL of supernatant was collected after centrifugation of the suspension for 10 min at 17,000×g and replaced with the same volume of fresh buffer solution. After the last washing and removal of the supernatant, precipitates were destroyed by a sodium citrate buffer (pH 5.5) at 40°C. The concentration of MB was determined by OD measurement using CLARIOstar PLUS plate reader.

### *In vitro* Analysis of Photodynamic Effect of MB-loaded NanoMOFs

Photodynamic effect of MB-loaded nanoMOFs and other formulations was measured using 2’,7’-dichlorofluorescin diacetate (H_2_DCF-DA) (Sigma-Aldrich, St Louis, MO), which converts upon ROS generation into fluorescent oxidation product. A stock solution of 4 mM H_2_DCF-DA was initially prepared in DMSO and then diluted to 5 µM with PBS. Then, H_2_DCF-DA (5 μM) was added to each of four samples: (1) pure nanoMOFs (100 µg mL^-1^), (2) MB-loaded nanoparticles (100 µg mL^-1^), (3) free MB (2 µg mL^-1^), and (4) a mixture of bare nanoMOFs (100 µg mL^-1^) and free MB (2 µg mL^-1^). Then, the samples were plated to 96-well plate and immediately irradiated with a diode laser (HTPOW, Seattle, WA) (λ = 650 ± 10 nm, output power 1 W) for 204 seconds (120 J cm^-2^). Fluorescent intensity of H_2_DCF-DA was measured using microplate reader CLARIOstar Plus after 1, 3, and 5 hours of incubation at 37℃ in the dark. As a control, we measured the same samples in a 96 -well plate without light irradiation.

### Cell Viability Assay

To estimate the cell viability, RAW264.7 cells were seeded in 96-well plate and incubated overnight at 5 % CO_2_ and 37°С before exposure to different concentrations (0, 5, 10, 50, 100, 250 μg per mL) of nanoMOFs, with or without MB, for 24 hours. A part of wells was irradiated upon 30 min incubation with formulations using a diode laser (HTPOW, Seattle, WA) (λ = 650 ± 10 nm, output power 1 W) at a dose rate of 120 J cm^-2^. CellTiter-Glo^®^ 3D Cell viability kit (Promega, Madison, WI) was used according to the manufacturer’s protocol.

### Evaluation of Antibacterial Effect in Infected RAW264.7 Macrophages upon Light Exposure

RAW264.7 cells were seeded into the wells of a 96-well plate with coverslips and cultivated for 24 hours at 37°C, 5 % CO_2_ prior to infection. The cells were infected with *C. trachomatis* strain L2/434/Bu at 100 MOI in SPG buffer for 2 hours by centrifugation at 500×g, followed by an additional incubation for 2 hours in standard conditions. Then, nanoMOFs, loaded or not with MB, or non-encapsulated MB were added to RAW264.7 cells at a final concentration of 100 µg mL^-1^ for MOF nanoparticles and 2 µg mL^-1^ for MB. After 30 minutes the infected cells were irradiated at a dose rate of 120 J cm^-2^ using a diode laser (HTPOW, Seattle, WA) (λ = 650 ± 10 nm, output power 1 W). After 48 hours of incubation, RAW264.7 cells were fixed with methanol, permeabilized with 1 % Triton X-100, and stained with ethidium bromide dye and FITC-labeled antibodies against chlamydial lipooligosaccharide (Galart Diagnosticum, Russia). Another 96-well plate with RAW264.7 cells was frozen at -70⁰C to obtain cell lysates. RAW264.7 cell lysates were collected, centrifuged for 10 min at 1,000×g and 4°C to get rid of cell debris, diluted in DMEM growth medium, and transferred to HeLa cells grown on coverslips in 24-well plates. Upon 48 hours of incubation, Hela cells were also fixed, and ICC stained in the same manner. All samples were prepared in triplicates. Images of 6 randomly selected fields from each stained coverslip (totally 18 fields per sample) were analyzed using the confocal microscope Zeiss LSM 710 (Carl Zeiss, Oberkochen, Germany) equipped with 20x/0.5 air objective lens. Chlamydia virulence was determined by quantification of fluorescent inclusions in 18 random fields of view. In addition, the infection rate was determined as the area of inclusions normalized to the area of HeLa cells. The images of the cells were quantified using Fiji software (version 2.1.0/1.53c).

## Acknowledgements

The authors acknowledge the Ministry of Science and Higher Education of the Russian Federation for financial support (Agreement № 075-15-2021-1363). Zhihao Yu acknowledges the China Scholarship Council for its financial support (CSC grant number 202008440250). The authors are also thankful to Dr. Andrey Bogorodskiy (Moscow Institute of Physics and Technology) for help with confocal microscopy experiments, Dr. Farid Nouar (Institute of Porous Materials from Paris) for help with preparation of figures, and Mr. Thomas McClymont (Institute of Porous Materials from Paris) for proof-reading of the manuscript. A part of cell culture experiments was performed using the facilities provided by the Center for Precision Genome Editing and Genetic Technologies for Biomedicine and Federal Research and Clinical Center of Physical-Chemical Medicine of Federal Medical Biological Agency.

## Author contributions

D.M., S.C., L.V., L.M. and Z.N. conceived and supervised the work; E.M. and M.A. performed the TEM measurements and data analysis; Q.X. and Y.Z. produced MB-loaded MIL-100 nanoparticles and characterized them; S.N. assisted in the experiments. G.E. and F.E. performed the experiments with *C. trachomatis*; Q.X., S.N., Y.Z., M.A. and M.D. prepared figures; Q.X., S.N. and Y.Z. wrote the manuscript with support of D.M., S.C. and L.M.; all authors contributed to the general discussion.

## Supporting Information

The Supporting Information is available free of charge at …

