## Supporting Information for "NanoMOFs with Encapsulated Photosensitizer: Accumulation in *Chlamydia trachomatis* Inclusions and Antimicrobial Effects"

**Table S1.** Physicochemical characteristics of MIL-100(Fe) nanoMOFs

|  | Mean diameter (nm)* | Mean diameter (nm)** | | PdI** | Zeta potential (mV)** | |
| --- | --- | --- | --- | --- | --- | --- |
| MIL-100(Fe) | 57 ± 10 | 147 ± 5 | 0.12 ± 0.01 | | | -29.7 ± 0.8 |

* Measured by SEM

** Measured by DLS

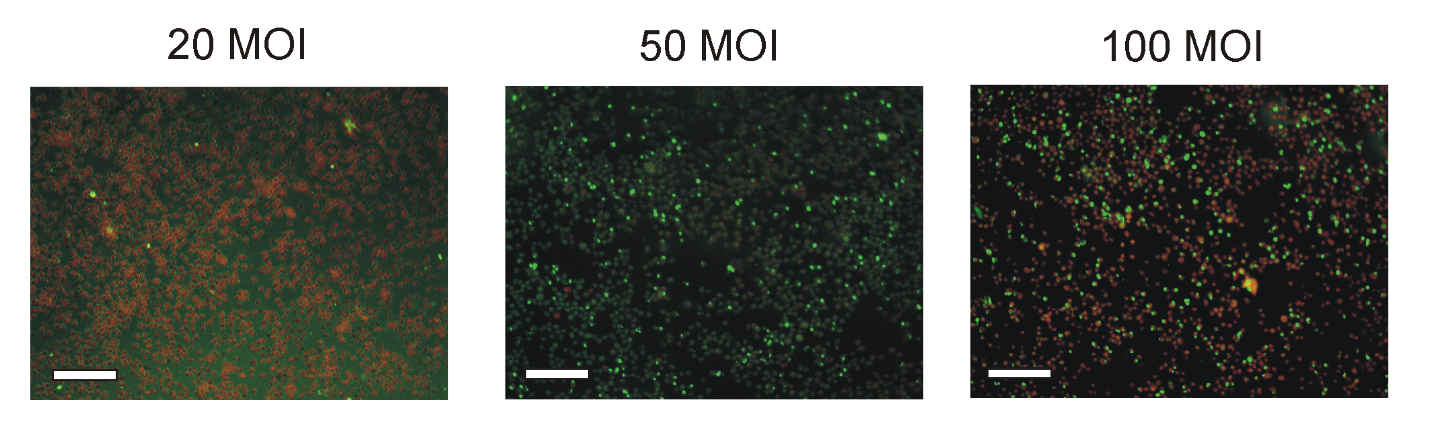

**Figure S1.** Selection of an optimal infection dose for RAW264.7 macrophages. After 48 hours of incubation with 20, 50 or 100 MOI, RAW264.7 macrophages were stained with Evans blue to visualize the cells (red) and FITC-labeled antibodies against chlamydial major outer membrane protein to visualize inclusions (green). Scale bar is 50 µm.

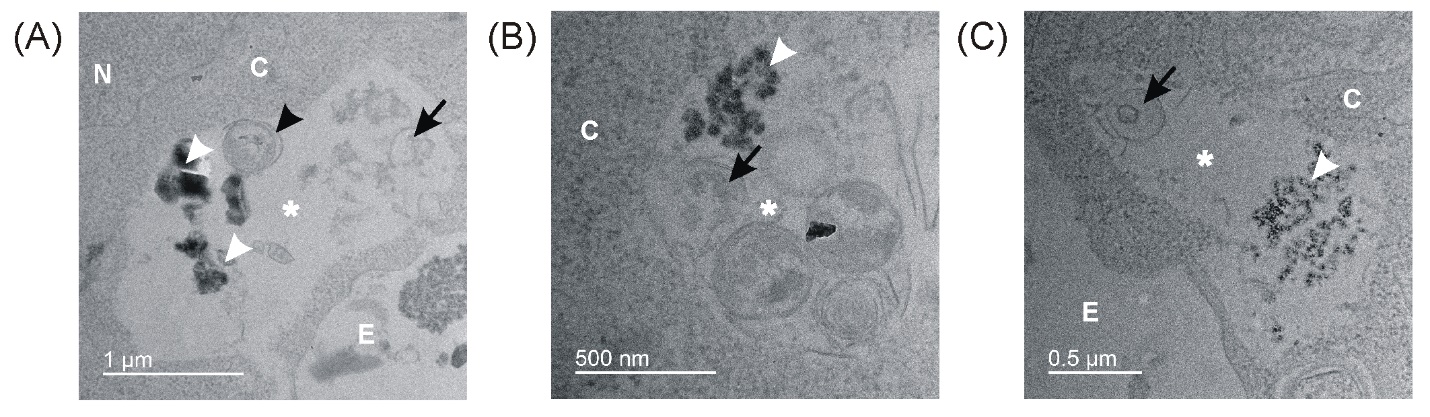

**Figure S2.** Co-localization of MIL-100(Fe) nanoMOFs with *C. trachomatis* ERs within inclusions in RAW264.7 cells. (a)-(c) Black and white arrowheads indicate chlamydial ERs and nanoMOFs, respectively. Black arrows indicate damaged bacteria. White stars indicate chlamydial inclusions. Abbreviations: N, nucleus; C, cytosol; P, phagosome; E, extracellular space.

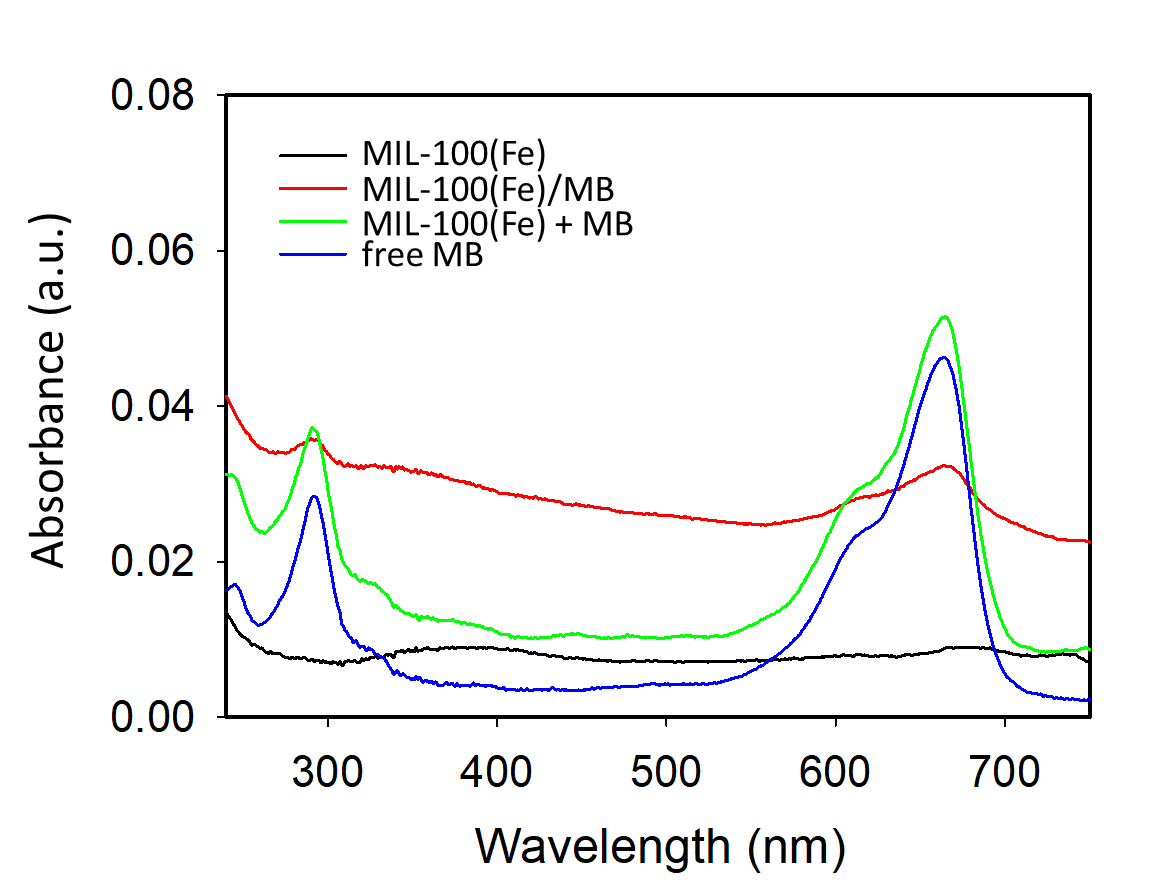

**Figure S3.** UV-VIS spectral absorbance of MIL-100(Fe) and MB formulations.

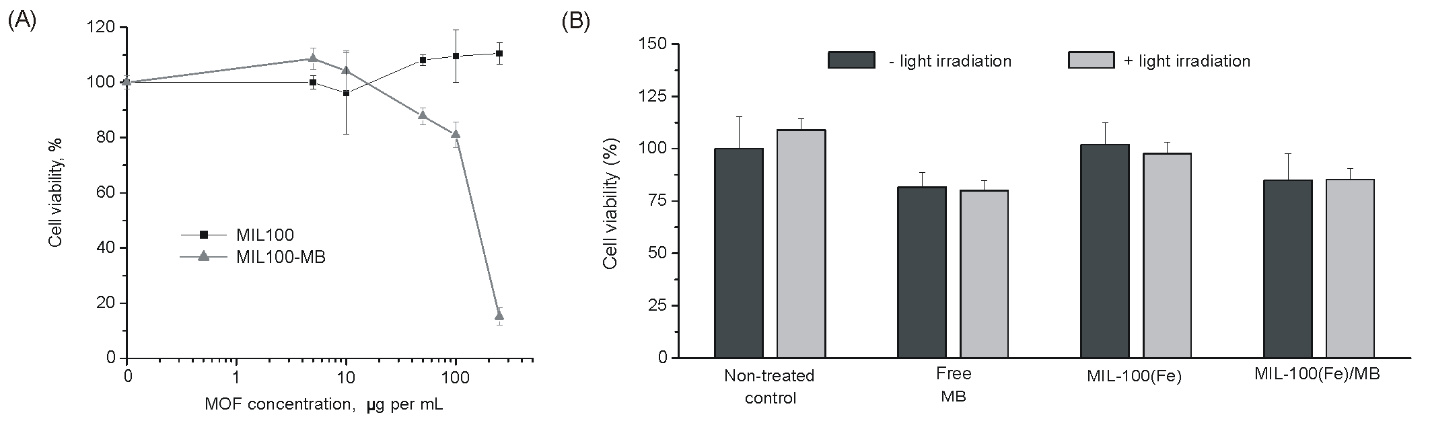

**Figure S4.** Analysis of cell viability after treatment with MIL-100(Fe) and MIL-100(Fe)/MB nanoMOFs. (A) Viability of RAW264.7 cells after 24-hour dark incubation with different concentrations of MIL-100(Fe) and MIL-100(Fe)/MB nanoMOFs. (B) Measurement of RAW264.7 cell viability after incubation with MIL-100(Fe), MIL-100(Fe)/MB nanoMOFs, or non-encapsulated MB, which were added at final concentrations of 100 µg mL^-1^ for MOFs and 2 µg mL^-1^ for MB. The cells were irradiated or not irradiated with light at a dose rate of 120 J cm^2^ and incubated for 48 hours. The data are given as means±SD.

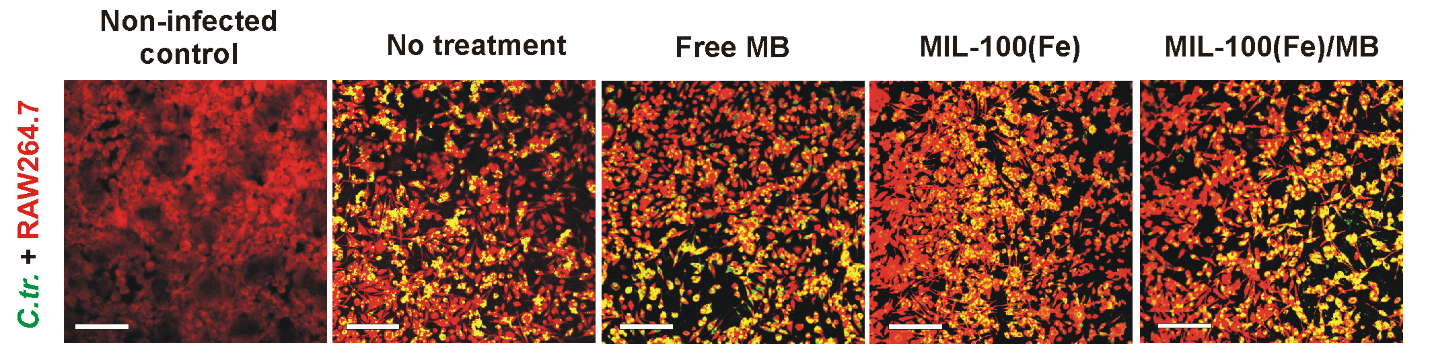

**Figure S5.** Images of infected RAW264.7 cells treated with MIL-100(Fe)/MB nanoMOFs and other formulations, followed by light irradiation. After 48 hours of incubation RAW264.7 macrophages were stained with Evans blue to visualize the cells (red) and with FITC-labeled antibodies against chlamydial lipooligosaccharide to visualize chlamydial inclusions (green). Scale bar is 50 µm.
